# Impact of *Itga2-Gp6*-double collagen receptor deficient mice for bone marrow megakaryocytes and platelets

**DOI:** 10.1101/625038

**Authors:** Daniela Semeniak, Kristina Faber, Patricia Öftering, Georgi Manukjan, Harald Schulze

## Abstract

The two main collagen receptors on platelets, GPVI and integrin α2β1, play an important role for the recognition of exposed collagen at sites of vessel injury, which leads to platelet activation and subsequently stable thrombus formation. Both receptors are already expressed on megakaryocytes, the platelet forming cells within the bone marrow. Megakaryocytes are in permanent contact with collagen filaments in the marrow cavity and at the basal lamina of sinusoids without obvious preactivation. The role of both collagen receptors for megakaryocyte maturation and thrombopoiesis is still poorly understood. To investigate the function of both collagen receptors, we generated mice that are double deficient for *Gp6* and *Itga2*. Flow cytometric analyses revealed that the deficiency of both receptors had no impact on platelet number and the expected lack in GPVI responsiveness. Integrin activation and degranulation ability was comparable to wildtype mice. By immunofluorescence microscopy, we could demonstrate that double-deficient megakaryocytes were overall normally distributed within the bone marrow. We found megakaryocyte count and size to be normal, the localization within the bone marrow, the degree of maturation, as well as their association to sinusoids were also unaltered. However, the contact of megakaryocytes to collagen type I filaments was decreased at sinusoids compared to wildtype mice, while the interaction to type IV collagen was unaffected. Our results imply that GPVI and α2β1 have no influence on the localization of megakaryocytes within the bone marrow, their association to the sinusoids or their maturation. The decreased contact of megakaryocytes to collagen type I might at least partially explain the unaltered platelet phenotype in these mice, since proplatelet formation is mediated by these receptors and their interaction to collagen. It is rather likely that other compensatory signaling pathways and receptors play a role that needs to be elucidated.

## INTRODUCTION

Blood platelets are key regulators to maintain vascular integrity and seal wounds after vessel injury. Recognition of subendothelial matrix proteins like collagen that become exposed after vessel injury is a trigger to bind and activate platelets, allow adhesion and aggregation and finally mediate thrombus formation under distinct flow and shear conditions. Platelets are derived from bone marrow (BM) megakaryocytes (MKs) that can transform their cytoplasmic content at the endothelial barrier of the microvasculature, where they shed proplatelets and platelets into the blood stream [1]. Within the BM, MKs are in permanent contact with proteins of the extracellular matrix, especially collagens, fibronectins, laminins or cross-linking proteins like perlecan or nidogen [2, 3]. Several collagen receptors are expressed during MK differentiation from hematopoietic stem and progenitor cells to mature MKs and finally platelets. LAIR-1 and DDR-1 are expressed on early precursors and for human cells co-expressed with CD34 [4, 5]. During MK maturation these markers are downregulated and concomitantly GPVI and the α2 integrin become upregulated. Circulating platelets express only these two receptors on their cell surface. Many assay systems have been developed to study the role of both receptors for adhesion under static and flow conditions as well as for thrombus formation in several systems focusing on different shear rates. Platelet adhesion on collagen under high shear does not require the two integrin receptors α2β1 and αIIbβ3 [6]. Platelet-collagen adhesion under flow, however, depends on the presence of α2β1 [7], but not for platelet-platelet aggregate formation on adherent platelets. The α2β1 integrin is not required for thrombi formation [8].

Both, GPVI and α2β1 have thus non-redundant functions and our current understanding states that GPVI is the central receptor for collagen on platelets [9]. Of note, integrin α2 still harbors non-redundant functions under certain experimental settings [10]. As antibodies directed against GPVI (like JAQ1) can deplete the receptor from the surface of circulating platelets without affecting the peripheral platelet count, this system has been used in parallel to the genetic ablation of α2β1, allowing to generate mice that are double-deficient for both receptors [11].

GPVI is a type I transmembrane receptor with a short cytoplasmic tail providing a docking site for Src family kinases (SFK). It is associated with the FcRγ chain which contains a classical immunoreceptor tyrosine-based activation motif (ITAM). Phosphorylation on tyrosine residues within this motif mediates Syk binding and finally activation of the LAT signalosome and PLCγ2 activation [12]. GPVI-deficient platelets have been generated and studied extensively [13]. On mouse platelets, loss of the FcRγ chain leads to concomitant lack of GPVI on the cell surface and *FCER1G*-null mice have been used as an animal model for GPVI-deficiency [14]. Of note, FcRγ-deficient animals might not fully reflect the *GP6*-null phenotype [10].

LAIR-1 is expressed on nearly all immune cells as well as on early hematopoietic stem and progenitor cells. In humans, it becomes downregulated concomitantly with CD34, when the stem cells differentiate into the megakaryocytic lineage [15–17]. LAIR-1 is a type I receptor with two immunoreceptor tyrosine-based inhibitory motifs (ITIM)s in its cytoplasmic domain. LAIR-1 binds to the collagen GPO motif with higher affinity than GPVI and can - when co-expressed - counteract the stimulatory effect of GPVI [18]. Mice lacking LAIR-1 show a defect in MKs, harbor a mild thrombocytosis and their platelets are surprisingly hyper-responsive, although LAIR-1 itself is undetectable in platelets. The underlying mechanisms have not yet been elucidated [4].

Several groups studied the effect of extracellular matrix proteins on proplatelet formation, mostly indicating that collagen type I is a negative regulator, while collagen type IV supports proplatelet formation. Recently, we developed mixed substrate assays to delineate which subtype is dominant and how this signal is transduced. Our data strongly support that collagen type I (and type III) transduce an inhibitory signal via GPVI [3]. As *Gp6*-null animals do not show any premature or ectopic proplatelet formation into the marrow cavity, we suggest that other signals besides GPVI contribute to the inhibition and provide part of a composite brake to prevent ectopic proplatelet release.

In this report, we aimed to study megakaryocyte maturation and thrombopoiesis in animals lacking both late collagen receptors GPVI and α2β1. Our results imply that MK maturation, their localization within the bone marrow or their association to the sinusoids is mediated by these receptors but both receptors are dispensable for the distribution of MKs within the bone marrow and for platelet formation.

## MATERIALS AND METHODS

### Animal Husbandry

Mice were housed in cages (max. 5 mice per cage) containing sawdust bedding and a nest made from paper tissues. The cages were kept at 23°C and underwent a 12-hour light phase followed by a 12-hour dark phase. Mice were maintained following the guidelines for animal care and welfare according to Tierschutz-Versuchstierverordnung with the approval of the district government of Lower Franconia (Bezirksregierung Unterfranken), in accordance with guidelines of the European Union. Mice were sacrificed by cervical dislocation under isofluran anesthesia. For this method, a general ethics waiver is granted by the Institutional Animal Care and Use Committee (IACUC) of the University Würzburg. Euthanasia was performed in a separate area away from other animals and all efforts were made to minimize suffering. *Gp6*-null and *Itga2*-null mice were used as recently described [3].

### Flow cytometry

Expression levels of platelet surface receptors, activation of αIIbβ3 integrin (CD41/61) or P-selectin exposure (CD62P) upon platelet stimulation with different agonists was determined by bleeding mice (100 μl) from the retrobulbar plexus of isoflurane-anesthetized mice into heparin filled reaction tubes. 1 ml of Hepes-Tyrodes buffer (HTB) was added and 50 μl of diluted blood was given to 10 μl of the respective fluorophore-conjugated antibody and incubated for 15 min in the dark. The reaction was stopped by adding 500 μl PBS and analyses were performed at a FACSCalibur flow cytometer using Cell Quest software (BD Biosciences). For the measurement of integrin activation and granule secretion, the remaining blood was washed twice with HTB. 50 μl of washed platelets were incubated with 7 μl agonist (final concentrations: 3 μM U46619; 10 μM ADP; 0.001 to 0.1 U/mL thrombin, or 0.1 to 10 μg/mL collagen related peptide [CRP-X_L_]) for 6 min at 37°C and 6 min at room temperature (RT) in a flow cytometry tube containing fluorophore-conjugated anti-αIIbβ3 and anti-P-selectin antibodies. By adding 500 μl PBS the reaction was stopped and analyses performed as described above.

For dense granule degranulation analyses, mice were bled 1:10 in acidic citrated dextrane (ACD), 25 μM mepacrine was added and samples were incubated for 30 min in the dark at 37°C. The reaction was stopped with 1.5 ml HTB. Measurement was performed at a FACSCelesta flow cytometer with FACS Diva software (vers. 8.0.1.1) by recording a mepacrine-unloaded sample for 60 sec followed by analysis of the mepacrine-loaded sample for 60 sec. Afterwards, degranulation was triggered by 0.1 U/ml thrombin and the decline of the mepacrine-based fluorescence signal was recorded for 180 sec indicating delta granule release.

MK ploidy analysis was performed on a FACSCelesta flow cytometer. Bone marrow was flushed out of femura and tibia bones from four mice, unspecific binding sites were blocked with an anti-Fc receptor antibody (2.4G2) for 20 min and MKs stained for additional 20 min with an anti-αIIbβ3 antibody. Samples were washed, fixed with 0.5% paraformaldehyde (PFA) and permeabilized with 0.1% Tween. DNA was stained with propidium iodide (PI) solution containing RNase over night at 4°C in the dark.

### Blood parameter

For platelet count and size as well as other blood parameters two drops of blood were taken from the retrobulbar plexus of isoflurane-anesthetized mice and analyzed by a scilVetabcPlus^+^ hemacytometer (Horiba materials).

### Immunofluorescence

Mice were sacrificed and femora and tibia isolated. Bones were fixed in 4% PFA for 4 h at 4°C prior and transferred to a serial gradient of sucrose solutions of 10, 20 and 30% (w/v), each for 24 h at 4°C and finally embedded in SCEM medium into cryo mold dishes, deep-frozen and kept at −80°C until sectioning. For IF-staining, organs were sliced into 10 μm sections on Kawamoto adhesive films with a cryotom (Leica CM1900) and fixed on superfrost glass slides as described recently [3]. For IF-staining, sections were thawed and rehydrated for 20 min in phosphate-buffered saline (PBS). Unspecific binding was prevented using a blocking buffer, sections were incubated with primary antibodies (GPVI and α2β1 from Emfret; Eibelstadt, Germany); CD105 from eBioscience (San Diego, CA, USA); collagen I from Abcam (Cambridge, UK), or collagen IV from Merck Millipore (Darmstadt, Germany) for 45 min at RT. Slides were washed three times for 5 min with PBS containing 5% fetal bovine serum (FBS) and 0.1% Tween20 and incubated with the respective secondary antibodies (anti-rat IgG Alexa 546 and 647; anti-rabbit IgG Alexa 647 and Alexa 568, or anti-goat IgG Alexa 647; all Life Technologies Carlsbad, CA, USA) for 45 min at RT. Blocking and antibody-incubation steps were repeated for each protein of interest. Nuclei were stained after drying the sections using DAPI-containing mounting medium Fluoroshield (Sigma-Aldrich; Schnelldorf, Germany).

Images were taken using a TCS SP8 confocal laser-scanning microscope (Leica Microsystems CMS, Wetzlar, Germany) with a 40x oil objective (NA 1.3) for a general map or a 63x oil objective (NA 1.4) with digital zoom for detailed records. Image documentation and analysis was performed with LAS X software, for MK measurements, images were processed with FIJI software [19].

### Data acquisition and statistical analyses

Statistical calculations were performed with GraphPad Prism (version 7, GrapPad Software, San Diego, USA). Results are shown as mean ± SD from three individual experiments, unless indicated otherwise. Significant differences of pairwise comparison of means against a specified control were determined using ANOVA, differences between cell size distributions of different groups were compared by Kolmogorov-Smirnov-test and frequency distributions by χ^2^-test. P-values <0.05 were considered as statistically significant with: p<0.05 (*), p<0.01 (**) and p<0.001 (***).

## RESULTS

*Gp6*-null and *Itga2*-null mice were present in our facility and interbred to allow littermate controls for single knockout (ko) lines as well as for the corresponding wildtype (wt) mice. Mice were intercrossed in a C57BL/6 background to minimize the effect of strain differences. Platelet counts and mean platelet volume (MPV) were unaffected in the double ko (dko) mice compared to single ko and wt mice (Fig 1A, B). Blood parameters were evaluated by two distinct blood cell counters and, despite some variations in few values (especially the white blood count), the red blood cell lineage and the leukocytes were overall unaffected in the dko mice (Fig 1C). Deletion of GPVI, α2 integrin or both collagen receptors on the platelet surface did not lead to an altered expression of von Willebrand receptor subunits GPIb, IX, V, the tetraspanin CD9, the integrin receptor αIIbβ3, or the hem-ITAM receptor CLEC-2 (Fig 1D). Absence of the collagen receptors was confirmed using antibodies directed against GPVI or α2 integrin and resulted in a nearly abrogated fluorescence signal in the respective blood samples. As shown for α2 integrin deficiency, we found compensatory upregulation of α5 integrin in mice lacking α2 integrin or in the dko platelets (Fig 1E). Taken together, our results clearly demonstrate that platelets from mice double deficient for both collagen receptors do not show any significant alterations in the surface receptor expression, with the exception of slight upregulation of α5 integrin subunit, which is attributed to the *Itga2*-null phenotype

**Fig 1.**
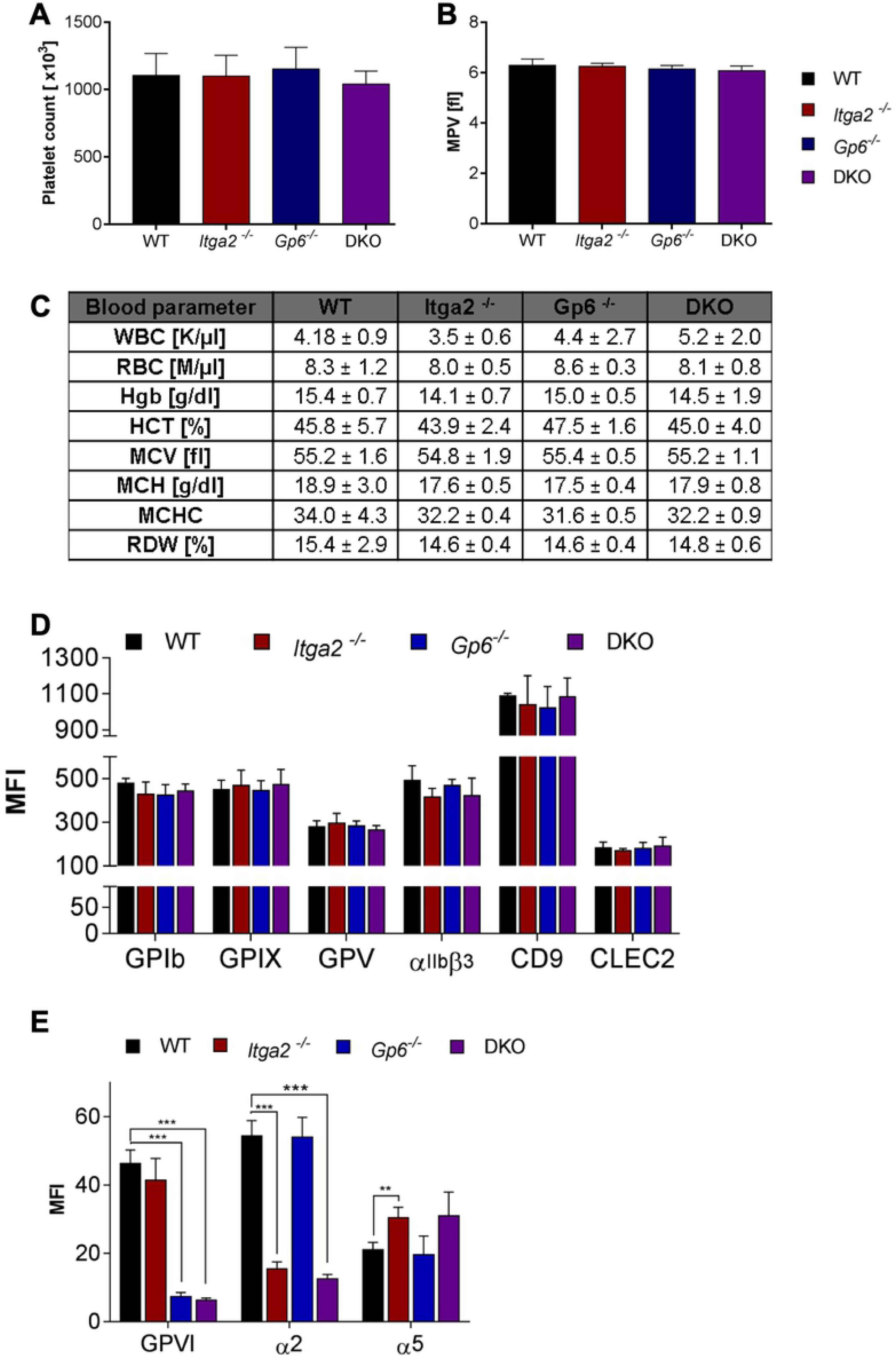
Platelet and blood parameter analysis in *Itga2/Gp6* double deficient mice. (A) Mice lacking collagen receptors GPVI and/or α2β1 integrin showed an unaltered platelet count and (B) size compared to wt controls. (C) Other blood parameters were also not influenced in dko mice. WBC = white blood cell concentration; RBC = red blood cell concentration; Hgb = hemoglobin; HCT = hematocrit; MCV = mean corpuscular volume; MCH = mean corpuscular hemoglobin; MCHC = mean corpuscular hemoglobin concentration; RDW = red blood cell distribution width. (D) No significant differences in receptor expression were found by flow cytometric analysis. (E) The absence of collagen receptors on the platelet surface was confirmed. Only the fibronectin receptor α5β1 showed higher MFI values in α2β1 single ko and dko compared to wt. Error bars indicate standard deviation. Asterisks mark statistically significant differences of the mean compared to wt mice (*** P < 0.001). Data of three independent experiments, each determined with 5 vs. 5 mice, were averaged.

Next, we studied platelet reactivity of dko platelets compared to the single ko platelets. Stimulation with 10 μM ADP alone or when ADP was combined with 3 μM of the thromboxane A2 receptor agonist U46619, integrin activation, monitored by binding of the epitope-specific antibody clone JON/A was unaffected (Fig 2A). We found that the dko platelets showed a slight hyperreactivity for alpha granule release in the ADP/U46619 co-stimulated platelets, as monitored by CD62P exposure, but the differences were statistically not significant (Fig 2B). When thrombin concentration was increased from 0.001 to 0.1 U/ml, we detected an unaltered reactivity for both integrin activation (binding of JON/A antibody) and for CD62P exposure, compared to single ko or wt platelets. As expected, the reactivity toward 1 or 10 μg/ml CRP-X_L_ was abrogated in GP6-null or in double deficient platelets. Finally, we asked whether dense granules were affected and used a mepacrine assay, where we monitored MFI values of unstained platelets, mepacrine-loaded platelets under resting conditions, and mepacrine-loaded platelets after stimulation with 0.1 U/ml thrombin for 180 seconds. We could not detect any difference between the three transgenic lines compared to the wt littermate controls (Fig 2C), suggesting that depletion of both collagen receptors does not lead to an overall change in platelet receptor expression, in reactivity to G-protein coupled receptors or in platelet granule composition or release.

**Fig 2.**
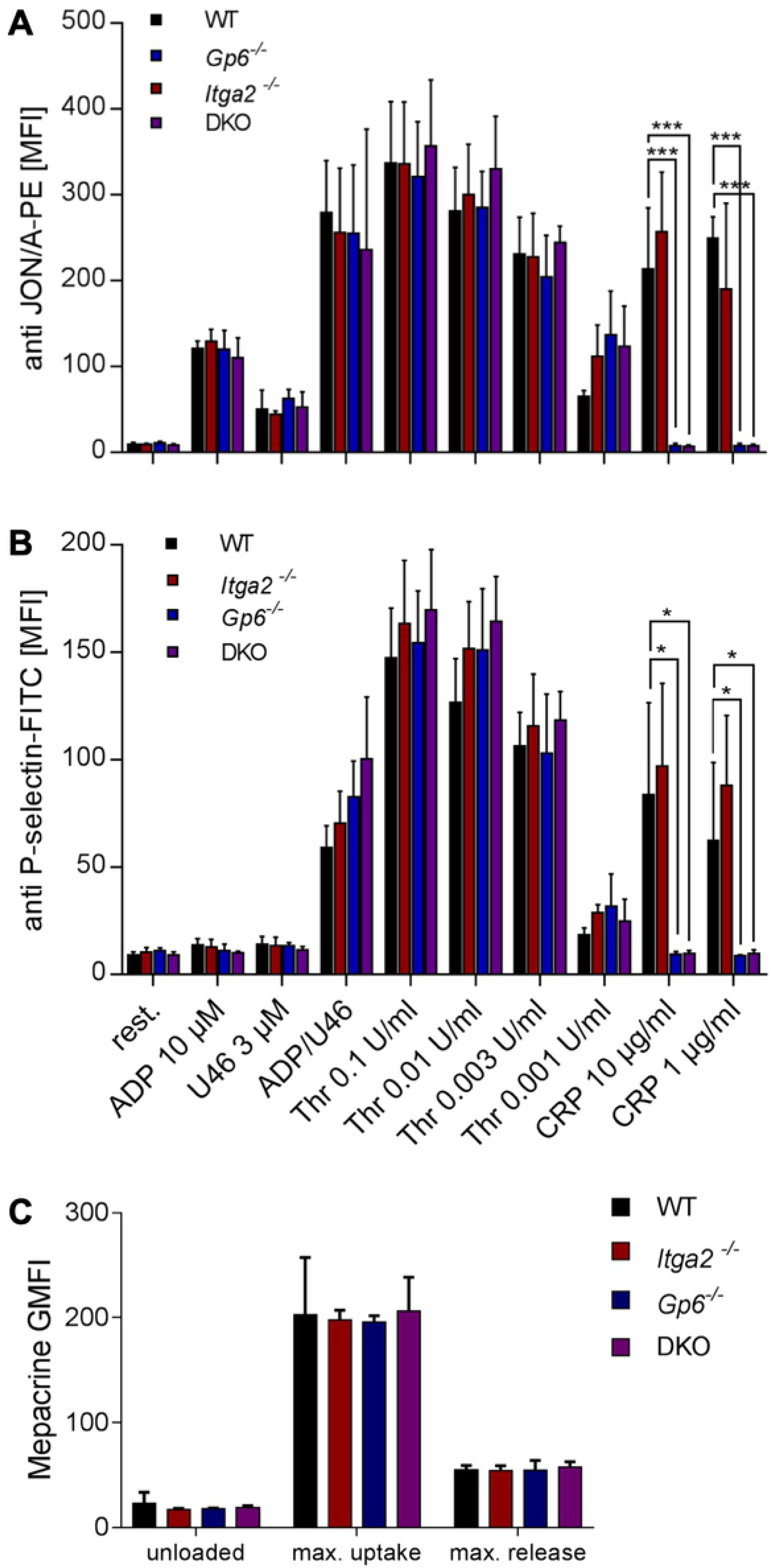
Ability of *Itga2*^−/−^/*Gp6*^−/−^ mice for integrin activation and degranulation. (A) αIIbβ3 activation was analyzed by binding of the epitope-specific JON/A-PE antibody, which showed the expected decrease of MFI in *Gp6*^−/−^ and dko mice upon CRP-X_L_ stimulation. (B) Platelet α-degranulation was measured by P-selectin exposure and *Gp6*^−/−^ and dko mice displayed decreased MFI values upon GPVI stimulation compared to wt. (C) The uptake and release of dense granule contents was measured by a mepacrine assay and alterations were present in single ko and dko compared to wt. Error bars indicate standard deviation. Asterisks mark statistically significant differences of the mean compared to wt mice (*** P < 0.001; * P < 0.05). Data of three independent experiments, each determined with 5 vs. 5 mice, were averaged.

So far, our experiments clearly state that none of both collagen receptors is required for platelet production. Moreover, the circulating platelets show an unaltered reactivity to all tested agonists with the exception of the GPVI-driven agonists CRP-X_L_ and convulxin. In order to exclude that the normal platelet count in double deficient mice was due to upregulation of bone marrow MKs, we studied their appearance in femur bones. Sections were fixed and subjected to specific antibodies. Nuclei were counterstained with DAPI prior to analysis by laser scanning-immunoflurescence microscopy. Both collagen receptors are expressed at low-copy numbers on MKs and we confirmed absence of GPVI, α2 integrin or both collagen receptors by co-staining sections with the respective antibodies, which resulted in the expected staining pattern: *Itga2*^−/−^ MKs showed no positive staining for α2 integrin and *Gp6*^−/−^ MKs no signal for GPVI, respectively. Dko MKs were negative for both proteins (Fig 3A). The unaltered concentration of circulating platelets in the peripheral blood was fully reflected by an unaltered MK number with about 12 MKs per visual field and the same MK area of about 320 μm^2^ (Fig 3B). MK ploidy was also unaltered in dko deficient animals as shown by PI staining (Fig 3C). These results suggest that MK maturation is unaltered in dko mice. Finally, we addressed whether lack of both collagen receptors would affect the localization within the bone marrow. To address this, femur sections were stained with anti-endoglin antibody (CD105), as a *bona fide* marker for bone marrow sinusoidal endothelial cells. The fraction of vessel-resident MKs was determined in comparison to marrow cavity-borne MKs. We found that almost half of MKs were in direct contact with vessels (Fig 3D), which is in agreement with previous studies [2, 3, 20] and that this is independent of the expression of both collagen receptors. Taken together, our results imply that both GPVI and α2 integrin are not required to generate MKs from hematopoietic stem cells under steady state conditions, that those MKs have an inconspicuous degree of maturation and that the MKs from double deficient animals localize to bone marrow sinusoids as their wt littermate controls.

**Fig 3.**
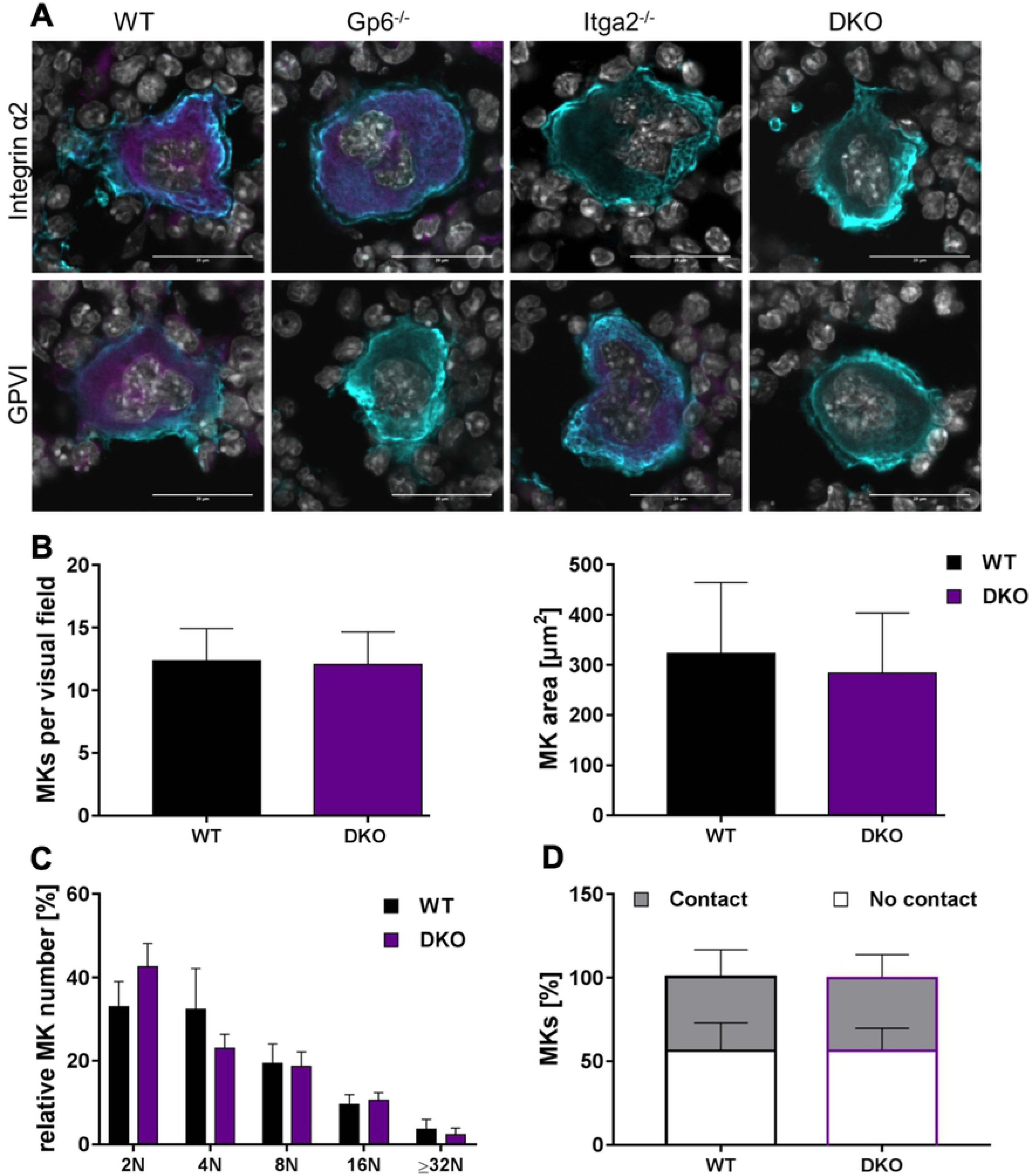
MK characterization and distribution within bone marrow of mice lacking the main collagen receptors GPVI and integrin α2. (A) Absence of GPVI and/or α2β1 (magenta) on the surface of MKs within bone marrow was confirmed by immunofluorescence staining of femur bone sections. MKs were identified by counterstaining GPIX (cyan). GPVI staining by JAQ1 antibody was not detectable in *Gp6*^−/−^ and dko MKs, as expected. Vice versa, no fluorescence signal was present in *Itga2*^−/−^ and dko mice when α2β1 was stained with LEN/B antibody. Nuclei were visualized with DAPI staining. Scale bars represent 20 μm. One of three representative experiments is shown. (B) Determination of MK number and size showed no differences between dko and wt controls. (C) Also the contact of MKs towards vessels and (D) the maturation state, measured by ploidy levels, were unaltered in dko and wt mice. Error bars indicate standard deviation. Data of three independent experiments, each determined with 3 vs. 3 mice, were averaged.

GPVI and the α2 integrin have been shown to mediate the adhesion of platelets or MKs to collagen filaments. Spreading on a collagen-coated surface was stronger for α2 integrin, rather than GPVI [3]. Thus, we asked next, whether lack of both receptors on megakaryoblasts and megakaryocytic progenitor cells leads to a change in association with collagen filaments. We thus stained femur sections with antibodies specific for either the collagen type I or type IV isoforms together with CD105 for the bone marrow endothelial cells and DAPI (Fig 4). We carefully characterized the used collagen antibodies and ensured that they are type-specific and do not show relevant cross-reactivity in respect to other collagen isoforms (for more details see also Fig S2, panels A-E in [3]). We could not detect any obvious differences when sections from double deficient animals were compared to the wt controls or in respect to the type of interaction with the sinusoids (Fig. 4A,B). Taken together, these results clearly suggest that the two collagen receptors are not required to guide maturing MKs within the bone marrow or to direct them toward the MK vascular niche (or the transendothelial passage).

**Fig 4.**
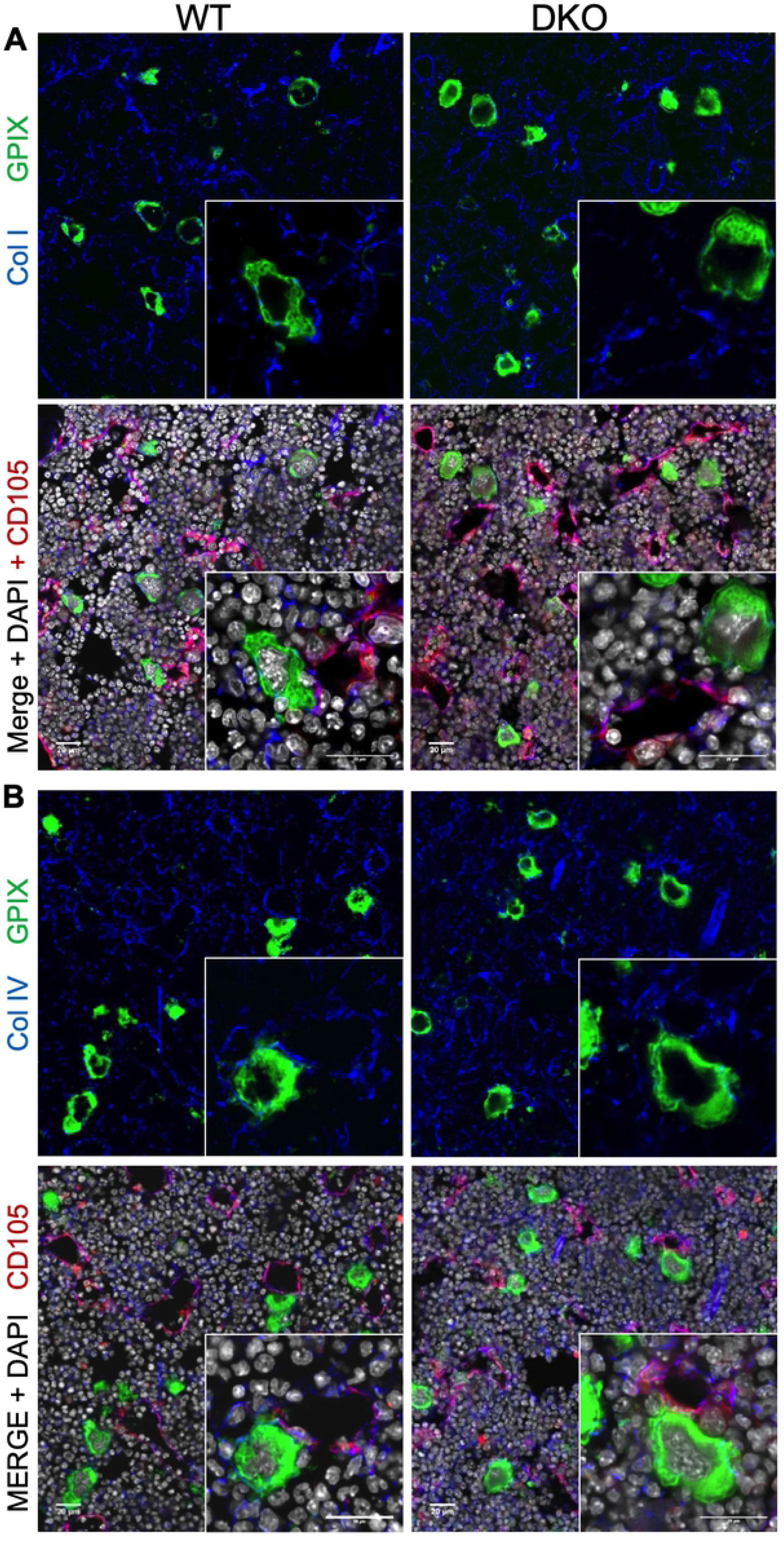
MK contact to collagens type I and IV within bone marrow of *Gp6/Itga2* double deficient mice. Collagen type I (A) or type IV (B) were stained together (blue) with endoglin (CD105), shown in red, and MKs (GPIX) in green. Nuclei were counterstained with DAPI (grey). Both collagen types are present at vessels and within the marrow cavity and enwrap MKs. No obvious differences in distribution or quantity of collagens were detectable between dko and wt mice. Scale bars represent 20 μm. One of three representative experiments is shown.

Finally, we aimed to address whether lack of both collagen receptors on MKs would affect the degree to which MKs are in direct contact with filaments from either collagen type I or type IV. For this approach, we decided to focus on MKs that have a rather low degree of interaction (less than 50%) and those MKs that show a significant contact with collagen filaments of 50% or more. Our analysis clearly revealed that MKs deficient for both collagen receptors lose contact to collagen type I. While about half of the wt MKs were in high contact with type I filaments, in dko mice only 30% showed a pattern of interaction, while 70% had only low contact (Fig 5A). This difference was statistically significant. However, when we co-stained for collagen type IV filaments, about half of MKs of either genotype were in high contact (Fig 5B), suggesting that the loss of collagen contact is specific for type IV.

**Fig 5.**
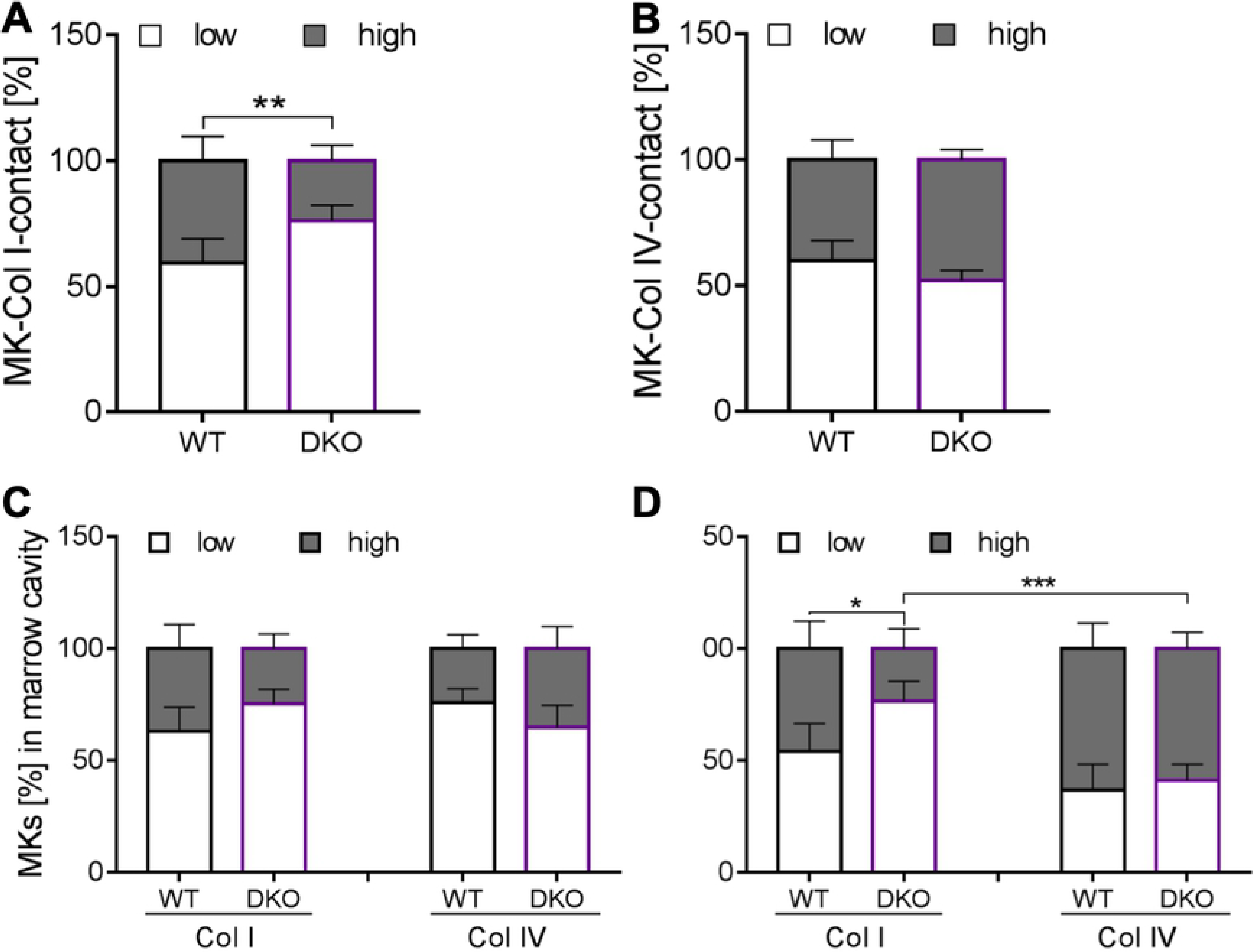
Degree of MK contact to collagen type I and IV within the marrow cavity and at the vascular niche. (A) The collagen type I contact to MKs was overall reduced in dko mice, whereas (B) Col IV contact was comparable to wt controls. While (C) in the marrow cavity the contact of collagen type I or type IV revealed no significant alterations between wt and dko mice, (D) at bone marrow sinusoids MKs had less contact to collagen type I, but demonstrated an unaltered contact to collagen type IV. Error bars indicate standard deviation. Asterisks mark statistically significant differences of the mean compared to WT controls (* P < 0.05; ** P < 0.01). At least 10 sections and 10 visual fields per section of 3 different mice were analyzed.

In order to exclude that this finding was confounded by the location of MKs within the bone marrow, we classified the degree of contact for every single MK in respect to those located in the marrow cavity and those that were vessel-associated MKs. When MKs were in the marrow cavity, there was no change in collagen interaction. We found a slight shift in regard to collagen type I over type IV in the absence of both collagen receptors (Fig 5C), but the effect was not statistically significant. In contrast, MKs of dko animals that were found to be at the vascular niche, were less densely packed with collagen type I, which reflects the finding described above. This difference becomes even more evident, when collagen types I and IV are directly compared for vessel-associated MKs: lack of both collagen receptors leads to MKs less densely associated with collagen type I compared to type IV (Fig 5D). These findings suggest that the expression of collagen receptors on MKs affects the direct interaction with collagen filaments and this association is more prominent for type I filaments compared to type IV filaments and becomes more evident at the vascular niche compared to MKs residing within the marrow cavity.

## DISCUSSION

The interaction of collagen with platelets has been broadly investigated over the last decades and key receptors and signaling pathways identified [9]. In contrast, the role of matrix proteins, like collagen, for MK signaling and proplatelet formation is much less well studied, with different approaches depending on the MK source (fetal liver, cord blood, bone marrow) and typically performed ex vivo and in vitro [21, 22]. Intensive studies about the interaction surface between MKs, matrix proteins and bone marrow vessels have become more readily feasible with the availability of new bone cutting and staining methods [23], more subtype-specific antibodies, as well as by the advent of novel microscopic approaches that allow to acquire both overview images of whole bones by maintaining high resolution images for detail analyses. This combination allows to deduce an organ-wide information on biological processes [24].

Thus, the main model of megakaryopoiesis and thrombopoiesis considers that hematopoietic stem cells, that reside in a quiet state at a vessel-distant side close to the bone (osteoblastic niche), do migrate within the marrow in response to stimuli and differentiate into terminally mature cells when they reach the endothelial blood-bone marrow barrier, where final platelet biogenesis occurs [25]. Recent data from complementing microscopy approaches have challenged this model with respect to limited bone surface in long bones in contrast to a dense vascular system that does not provide much space for migration [20]. While this model still has some shortcomings, it might provide an explanation of limited movement of MKs within the marrow cavity. This compartment contains a dense network of multiple matrix proteins, many of them expressed as spatially (or temporally) distinct isotypes with different to oppositional effects (as typically described for collagens type I and type IV). The inert reaction of MKs to collagens within the bone marrow has been a conundrum for many years.

While mice (and especially platelets) deficient for one or two collagen receptors have been addressed in multiple excellent studies, the role of collagen receptor-deficient MKs within the bone marrow has not yet been analyzed. In this report, we aim to address how HSC and megakaryocytic progenitor cells that are blind for direct collagen binding can find the vascular niche and release proplatelets and platelets.

### Ectopic platelet production into the marrow

All nucleated blood cells and red blood cells are released as (almost) mature cells into the circulation. Platelets, in contrast, are released as subcellular fragments from large precursor cells that function as a reservoir for membranes, granules and cytoskeletal proteins, allowing them to release up to 1000 mature platelets from a single MK [26]. The blood-bone marrow barrier is of special interest for this transition, as under healthy conditions we do not observe that platelets become prematurely (or ectopically) released into the bone marrow cavity rather than across the endothelial barrier into the blood stream. Some mice that lack regulatory proteins have been described to prematurely release platelet: WAS [27], CK2B [28], or ADAP [29]. The underlying mechanism, however, remains poorly described. Proplatelet formation can also be observed in vitro, especially from fetal liver-derived MKs [30], while bone marrow-derived MKs require some more attention like the addition of hirudin [31] or the use of explant models [32]. Our previous work has identified that the most common candidates for matrix proteins to modulate proplatelet formation in vivo (i.e. collagen type IV, laminin, fibronectin) have no stimulatory effect in vitro. In contrast, collagen type I (and to a lesser degree type III) have the power to inhibit (or to attenuate) proplatelet formation, which might explain why in the MK-collagen measurements only collagen type I association was impaired. This action is dominant over collagen type IV (or laminin) and mediated by collagen receptor GPVI [3]. As the α2 integrin is the major adhesive receptor for collagen, but did not affect the inhibition of proplatelet formation, we suggest a model where binding of type I and IV collagens differs between the marrow cavity and the vascular niche. 50 to 70% of mature MKs are found at the vessels, where they are enwrapped with type IV rather than type I filaments. This model allows to explain why proplatelet formation occurs at the vascular niche instead of into the marrow cavity. We suggest a model where proplatelet formation is set on hold with brakes on (to prohibit premature release), rather than awaiting a signal that triggers proplatelet formation. Obviously, the *Gp6*-null mouse does not show any premature proplatelets into the marrow, which can readily be explained by the presence of a second (or any additional) brake(s). The second collagen receptor integrin α2β1 is, of note, no second brake in this manner.

### Does co-localization mean “binding to collagen”?

The presented results for platelet function studies of double-deficient platelets are based on well-established assays. Our MK data, in contrast, is mainly based on images derived from confocal laser scanning microscopy. We used bone sections between 7 and 10 μm which allows to acquire a small stack of z-images. Classical bone sectioning protocols include a decalcification step, typically by using harsh conditions with concentrated hydrochloric acid. This procedure can destroy selective epitopes and thus lead to a biased mapping of extracellular matrix protein expression within the bone marrow. We adopted a modified bone cutting and staining protocol that does not require any decalcification. Our antibodies directed against different collagens are well-characterized and show a minimal overlap for the distinct types [3]. Many crosslinking proteins that can bind to several collagen types suggest that our overlap for filaments is quite realistic. Nonetheless, the general problem remains that we conclude a direct interaction (i.e. based on images from a collagen filament showing an elaborate wrapping pattern of a single MK) is somehow based on an interaction with collagen receptors present on the MK surface. We were to some degree surprised that MKs that do not express any of the collagen receptors still associate with (single) collagen filaments. Of note, we cannot exclude that other receptors on the MK surface that have collagen binding capacity on their own (i.e. GPV) mediate this interaction. Other β1-integrin family integrins might bind collagen indirectly due to interaction with their main ligand: MKs and platelets express the receptor for laminin (α6β1-integrin) or fibronectin (α5β1-integrin), the latter of which is known to be upregulated in *Itga2*-null and dko mice. Finally, the mere density of collagen filaments could result into the impression that MKs are in permanent contact with these filaments without any special receptor-based interaction. However, our data indicate that specifically interactions between MKs and collagen type I are altered when both receptors are missing (Fig. 5A,D), suggesting that the detected interaction is at least partially mediated by binding to collagen receptors.

### Other collagen receptors (compensatory mechanisms)

Our work has focused on the two collagen receptors that are expressed on both, platelets and megakaryocytes: the α2 integrin and GPVI. Early MKs do express other collagen receptors that are not expressed on platelets: DDR-1 expression has been detected on human MKs [5] and we could confirm the presence of DDR-1 on MKs in mouse femur sections (Fig. 6A). However, we detected a positive staining for DDR-1 mainly in GPIX-negative cells, independent of the presence or absence of collagen receptor expression. This makes DDR-1 an unlikely candidate for MK-collagen interactions in the bone marrow. LAIR-1 is a better characterized collagen receptor which belongs to the ITIM receptor family. Platelets of mice lacking LAIR-1 are still hyperreactive upon collagen stimulation, although the receptor is absent on platelets [4]. When we co-stained LAIR-1 together with MKs (GPIX) within bone marrow of wt and dko mice, we could detect single GPIX-negative cells that were strongly positive for LAIR-1, likely being early megakaryocytic progenitor cells. In addition, the more mature MKs showed a rather faint staining for LAIR-1. Nevertheless, the staining pattern was comparable between wt and dko animals, indicating that LAIR-1 is most likely not a compensatory collagen receptor in the compound absence of GPVI and α2 integrin (Fig. 6B). A second ITIM bearing receptor that might be of interest, is G6b-b. Although this receptor shares some signaling similarities with GPVI (phosphorylation of Src), it inhibits the phosphorylation of Syk and LAT downstream of Src family kinases. Full G6b-b knockout leads to a downregulation of GPVI [33], most likely due to receptor shedding.

**Fig 6.**
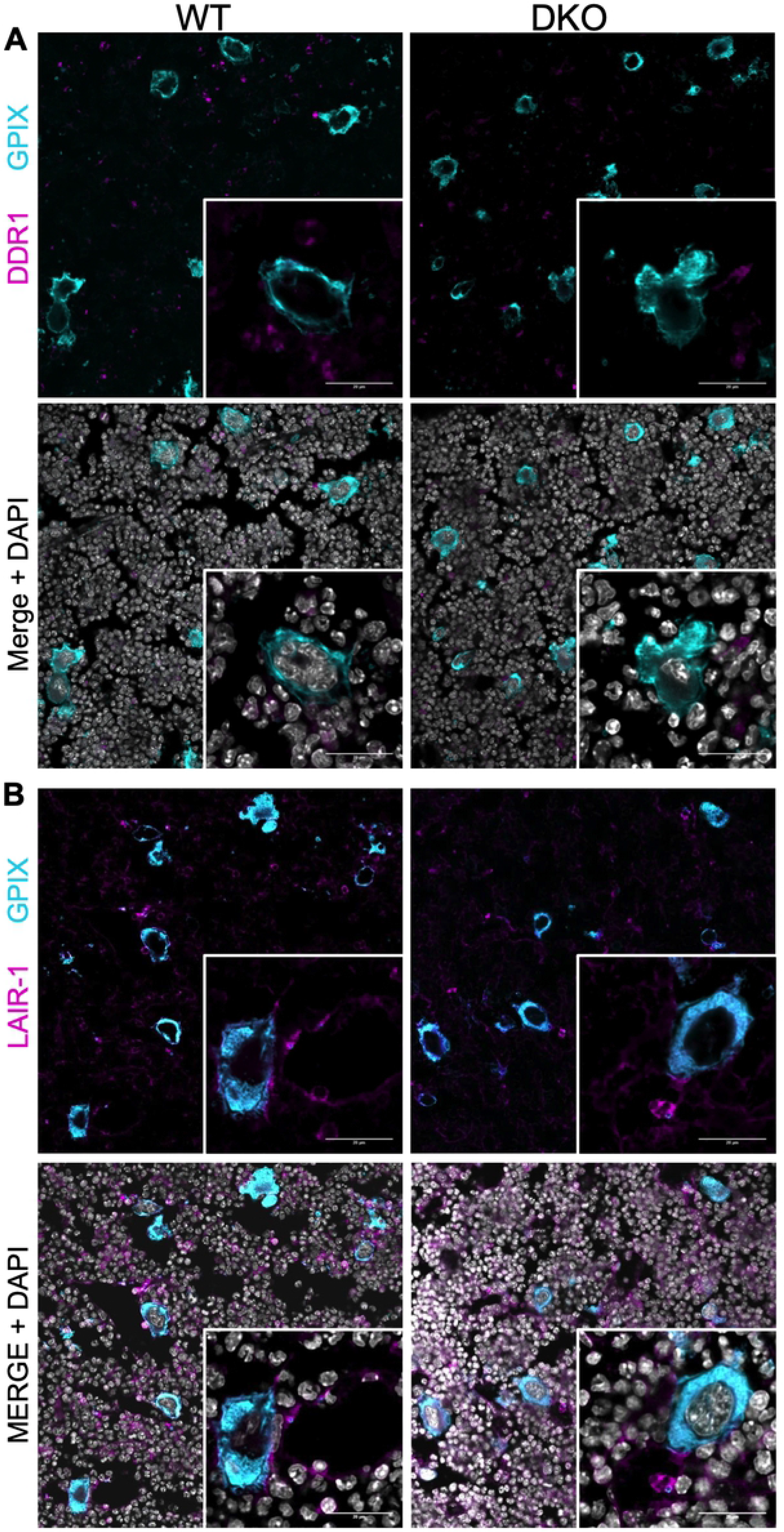
Expression pattern of early collagen receptors DDR-1 and LAIR-1 in mice double deficient for collagen receptors *Gp6* and *Itga2*. Either DDR-1 (A) or LAIR-1 (B) were stained together (magenta) with GPIX, a lineage marker for MKs (cyan). Nuclei were counterstained with DAPI (grey). (A) DDR-1 is expressed within bone marrow, but only in GPIX-negative cells. (B) Single bone marrow cells are markedly positive for LAIR-1, whereas all GPIX-positive MKs and sinusoids were only weakly positive. No obvious differences in distribution or quantity of these receptors were detectable between dko and wt mice. Scale bars represent 20 μm. One of three representative experiments is shown.

Another interesting observation has been reported in a study where platelets have been removed from the circulation by either anti-GPIIb/IIIa or anti-GPIb antibody application. Those platelets that are newly formed (on day 4-5 after depletion) do express GPVI on the cell surface, but this receptor cannot fully signal in response to GPVI agonists and downstream signaling molecules Syk and LAT do not become phosphorylated. The newly formed platelets failed to fully adhere on immobilized collagen in a flow chamber. The authors suggest that GPVI signaling is masked on newly released platelets and becomes activated when platelets are already in the circulation. This principle could also explain why bone marrow MKs do not become activated in response to collagen filaments [34].

Further hints for a compensatory or at least an additional signal would be the not totally abrogated proplatelet formation of MKs seeded on collagen type I. 10% of MKs were still able to form proplatelets on type I collagen, indicating that other receptors do contribute.

GPV can also bind collagen and *Gp5*-deficient platelets are unable to adhere to collagen type I under static and flow conditions [35], indicating the role of alternative binding options when both main collagen receptors GPVI and α2β1 are absent. The two main collagen receptors alone are thus dispensable for platelet production and not responsible for the prevention of ectopic platelet release. Our study is the first to use a genetic double knockout mouse model to study the effect of MK differentiation and maturation within bone marrow in respect to platelet biogenesis. The results clearly imply that MK-collagen interactions are dispensable for MK guidance within the marrow cavity or MK migration towards the blood vessels. This finding provides some insight that collagen receptor signaling differs substantially from peripheral platelets.

## ACKNOWLEDMENTS

The authors would like to thank the SFB688 (A21) and TR240 (A03) project for financial support, Nadine Winter for excellent technical support and Bernhard Nieswandt for critical comments on the manuscript.

